# Machine Learning Matches Human Performance at Segmenting the Human Visual Cortex

**DOI:** 10.1101/2025.05.16.654503

**Authors:** Noah C. Benson, Bogeng Song, Shaoling Chen, Toshikazu Miyata, Hiromasa Takemura, Jonathan Winawer

**Author notes:** Dept. of Psychology, Georgia Institute of Technology, Georgia, United States (Bogeng Song). Dept. of Psychology, Princeton University, Princeton, United States (Toshikazu Miyata).

## Abstract

A major problem in human visual neuroscience research is the localization of visual areas on the cortical surface. Currently available methods are capable of making detailed predictions about many areas, but human raters do not agree as well with these methods as they do with each other. Although highly accurate, human raters require substantial time and expertise that many researchers do not have. Additionally, human raters require functional data for drawing visual area boundaries that requires additional scan time, budget, and expertise to collect. Here, we train convolutional neural network (CNN) models to predict the boundaries and iso-eccentric regions of V1, V2, and V3 in both the Human Connectome Project dataset and the NYU Retinotopy dataset. CNNs trained to use the same functional data available to human raters predicted these boundaries with an accuracy similar to humans, while CNNs trained to use only anatomical data had a lower accuracy that was nonetheless higher than that of any currently available method. A comparison of the model accuracies when predicting eccentricity-based boundaries and polar angle-based boundaries suggests that eccentricity is substantially less closely tied to anatomical structure than polar angle and that the cortical magnification function, at least in terms of eccentricity, varies substantially between subjects. We further find that the fraction of V1, V2, and V3 that can be accurately parcellated into function regions using gray matter structural data alone is ∼75% (∼80% of the inter-rater reliability of human experts), implying a much tighter coupling between structure and function in these areas than previously estimated. We conclude that machine learning techniques such as CNNs provide a powerful tool for mapping the brain with human accuracy and predict that such tools will become integral to neuroscience research going forward.

## Introduction

The human visual cortex is tiled by many distinct visual areas whose functions are presumed to also be distinct. These visual areas begin with the primary visual cortex (V1), the first location on the cortex where information from the retina is processed, and immediately surrounding V1 are areas V2 and V3. These three regions are among the largest and most studied areas of the human brain and are of particular interest to researchers and clinicians who study blindness and degenerative visual disorders such as glaucoma, macular degeneration, and Leber hereditary optic neuropathy due to the fact that scotomata are generally reflected in these early visual areas (Baseler et al., 2011; Masuda et al., 2021). Both clinical research into such disorders and basic vision research frequently require that the investigators know the position and organization of these visual areas in their subjects in order to draw meaningful conclusions. However, the structure of these areas has enormous variability across subjects: V1’s cortical surface area can vary by at least a factor of 3, while the brain’s entire cortical surface area varies only by a factor of ∼1.5 (Benson et al., 2022; **Fig. 1**).

**Figure 1.**
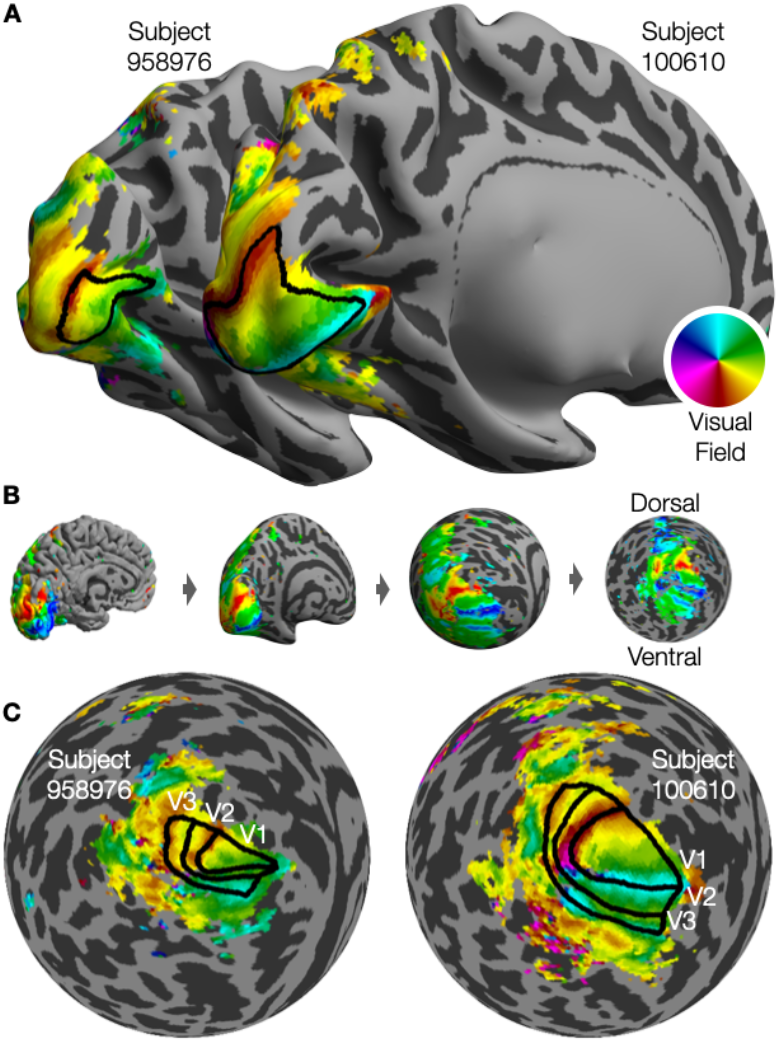
Visual areas are defined in terms of their retinotopic maps. (**A**) Two subjects from the HCP with polar angle maps and V1 boundaries (drawn by humans) shown. The boundaries of V1 correspond to the reversals in the polar angle map. (**B**) A schematic demonstrating how the cortical surface is inflated then flattened into a 2D orthographic projection. (**C**) Flattened polar angle maps for both subjects with V1, V2, and V3 boundaries shown.

Previous work into the early visual cortex found that, by aligning the cortical folding pattern of an individual subject to that of an anatomical template using FreeSurfer (Fischl et al., 1999), one can use a model of retinotopy defined in terms of the anatomical template to segment the approximate functional boundaries and organization of areas V1, V2, and V3 from anatomical information alone (Benson et al., 2014). If one additionally possesses functional magnetic resonance imaging (fMRI) measurements for the subject during a retinotopic mapping experiment (Sereno et al., 1995; DeYoe et al., 1996; Benson and Winawer, 2018), the predicted organization can be somewhat improved by using a high-dimensional optimization technique that aligns the measured retinotopic maps to those of the model of retinotopy (Benson and Winawer, 2018). Unfortunately, the accuracy of these predictions remains lower than that of visual area boundaries that are manually delineated by humans (Benson et al., 2022).

Computational problems such as image segmentation that are generally difficult for a traditional algorithm to solve but easy for a human to solve, can often be solved effectively using machine learning methods such as convolutional neural networks (CNNs), the design of which was originally inspired by the neural organization of the visual system itself (Lindsay, 2021). Previous work, in fact, has demonstrated that the retinotopic organization of the cortex can be accurately predicted by CNNs (Ribeiro et al., 2021). One particular class of CNN called a U-net has been developed specifically for medical image segmentation and for use with problems where the number of training examples is relatively small (Ronneberger et al., 2015; Siddique et al., 2021). Here, we employ a U-net to predict both the boundaries of visual areas V1, V2, and V3 and a set of orthogonal regions in V1, V2, and V3 delineated by iso-eccentricity curves in the 181 subjects of the Human Connectome Project 7 Tesla Retinotopy Dataset (Van Essen et al., 2013; Benson et al., 2018). These regions and their annotation were described in previous work (Benson et al., 2022). We find that CNNs trained to use structural information alone substantially outperform previous methods of predicting visual area boundaries, even those that employ functional data. Similarly, CNNs using structural data to predict regions delineated by eccentricity outperform previous methods that also use structural data alone but do not substantially improve the predictions of existing methods that exploit functional data. CNNs trained to use functional information as well as anatomical information, however, have an accuracy similar to that of humans for both visual area boundaries and boundaries delineated by eccentricity. Further, because these models offer the best predictions, to date, of V1–V3 organization, they provide new insight into the extent to which structure can be used to predict function in the visual cortex.

## Methods

This project involves the supervised training of several U-net CNNs to predict either the visual area boundaries of V1, V2, and V3 or a set of five regions delineated by eccentricity, which we refer to as iso-eccentric regions. Further, each CNN was trained to use a particular set of input data. All training was performed using disjoint training and validation sets, and hyperparameters were found using a grid search. Finally, models were validated against an independent dataset.

### Model Structure

We used an encoder-decoder based CNN structure to perform our image segmentation task. The details of the encoder model’s structure are described by (Ronneberger et al., 2015). In brief, the model contains two sections: the encoder section and the decoder section. The encoder is used to capture the features from the original images and encode them in a latent lower-dimensional space. After extracting the important features, the decoder is used to decompress these latent features by processing them through several steps that perform both convolutions and upsampling, eventually producing features that have been upsampled back into the size of the original input images. We chose a ResNet model, a well-known model architecture that has been previously studied for biomedical image classification (He et al., 2016; Xu et al., 2023), as our encoder model. The ResNet architecture contains several “residual blocks” used to capture detailed and hierarchical features that are beneficial for segmentation tasks. Each residual block contains convolution layers, batch normalization layers, and activation layers to extract useful information from the original input images. After data is processed by the ResNet encoder model into a latent encoding, that encoding is further processed by the decoder. The decoder component contains the same number of blocks as the encoder, which gradually reconstruct an image of the predicted visual area labels or iso-eccentric region labels. We use traditional bilinear up-sampling methods to expand the latent representation of the encoder back into our original input image size. Skip connection paths are additionally used in the U-net to transfer the outputs of each encoder block to the matched decoder block, preserving spatial information lost during encoding. Unlike traditional U-net models, our input images were not composed of RGB channels; instead we selected different fMRI data for each channel of our training inputs. Each channel of the input image can contain a different kind of data derived from the cortical surface, for example curvature or gray-matter thickness. All models used the same number of internal layers with the exception of the initial model layer, whose convolutional weights must match the number of channels in the input image. Accordingly, we adjusted the first layer in the encoder to match the number of channels in each model’s input images. We describe the input images in detail in the next section.

### Dataset

The fMRI data used in training were obtained from the Human Connectome Project (HCP; Van Essen et al., 2013), which contain a huge trove of neuroimaging data, including retinotopic mapping data from 181 subjects (Benson et al., 2018). The data acquisition and preprocessing details were described in previous publications.

Because our models require supervised learning, we used a set of hand-drawn labels that delineated both the V1, V2, and V3 boundaries for each subject as well as a set of orthogonal iso-eccentric regions. The iso-eccentric lines of delineation were drawn at 0.5°, 1°, 2°, 4°, and 7° forming 5 regions (0–0.5°, 0.5–1°, 1–2°, 2–4°, and 4–7°; **Fig. 2**). The visual area boundaries were also peripherally bounded at 7° of eccentricity. These regions were each drawn by 4 trained individuals using methods described previously (Benson et al., 2022), and each rater’s boundaries for each subject were used as a separate training example in our dataset.

**Figure 2.**
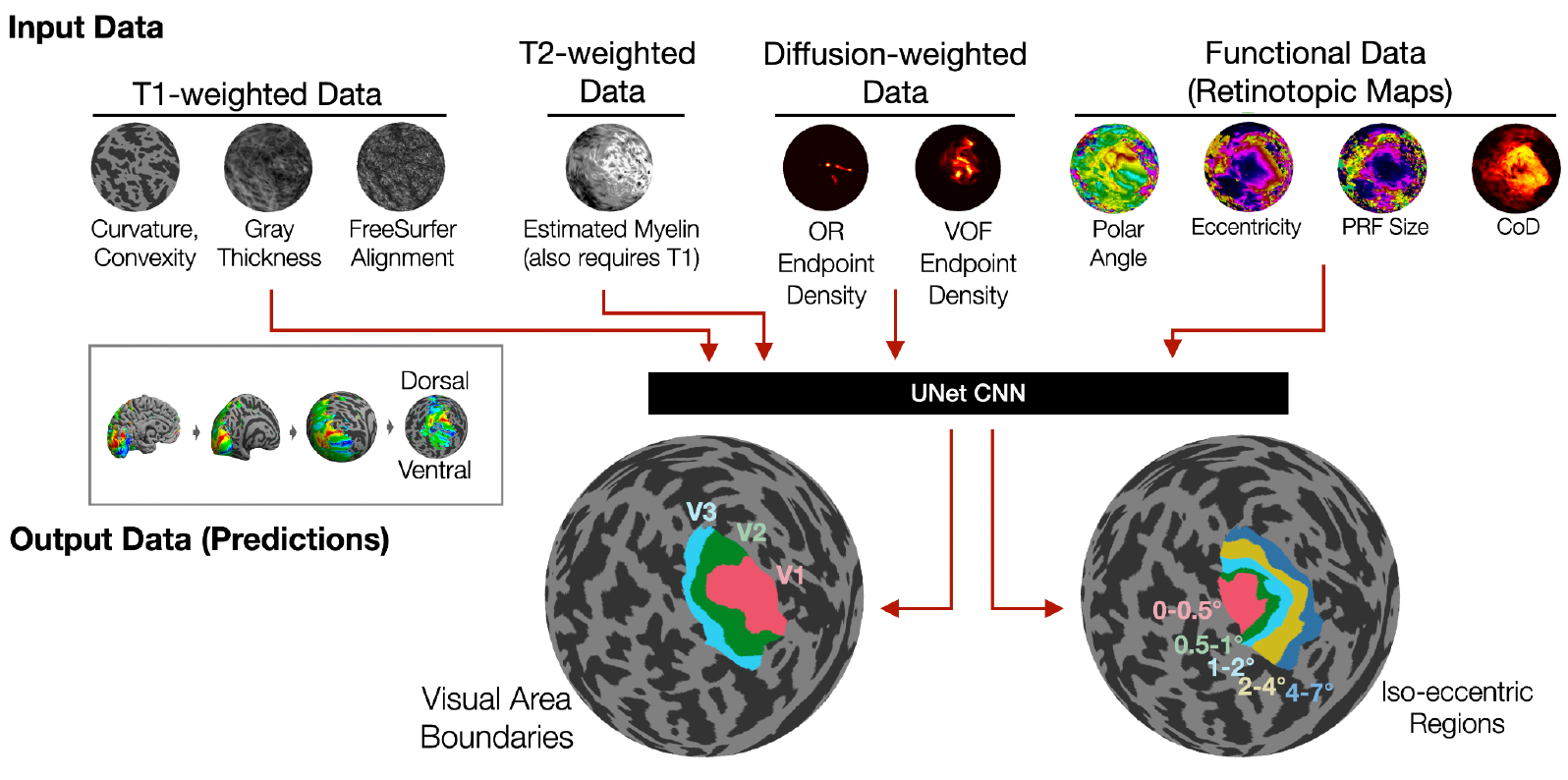
Overview of CNN training. The CNNs were trained using a variety of input data: (Group 1) T1-weighted data, (Group 2) T2-weighted data, (Group 3) diffusion-weighted data, and (Group 4) functional PRF mapping data. These data are used to predict either the subject’s V1, V2, and V3 boundaries or a set of 5 iso-eccentric regions orthogonal to the boundaries. A different CNN model was trained for every combination of input data from Groups 2, 3, and 4 (data from Group 1 was included in every CNN) and for each of the two outputs. For each Group except Group 1, the data were either included or excluded resulting in 2^3^ = 8 combinations of inputs and 2 outputs for 16 total CNNs trained. All input and output data were converted into 2D images by orthographic projection of the spherical *fsaverage*-aligned cortical surface.

To probe which fMRI data would be useful for segmenting the visual cortex, we used a variety of MRI data for each of the input image channels. We used four groups of images (**Fig. 2**). Group 1 included only anatomical data derived from T1-weighted images (curvature, convexity, gray-matter thickness, and vertex surface area, which implicitly includes information about the geometry of the surface and the FreeSurfer alignment); because a T1-weighted image is required in order to use the model, all data from Group 1 are also included in all other input groups. Group 2 includes anatomical data derived from T2-weighted images, specifically the HCP’s estimated myelin (T1w/T2w ratio map; Glasser and Van Essen, 2011). Group 3 contains white matter tract endpoints from diffusion-weighted data, specifically endpoint density maps for the Optic Radiation (OR) and Vertical Occipital Fasciculus (VOF), which are generated by a method described in previous work (Takemura et al., 2016). Finally, Group 4 includes functional data from retinotopic mapping, specifically the polar angle, eccentricity, pRpF size, and Coefficient of Determination (CoD) previously published by Benson et al. (2018). We trained models to use eight different combinations of these data groups: each input combination included Group 1 and either included or did not include each of Groups 2, 3, and 4. Each model additionally predicted either visual area boundaries or iso-eccentric regions, yielding 16 total models.

Input image channels were each produced from a single property (e.g., curvature or eccentricity; **Fig. 2**) by rendering orthographic projections of each subject’s left and right hemispheres side by side in a 128 × 256 pixel grayscale image. **Figure 2** shows examples of these orthographic projections, though for clarity only the left hemisphere is shown and some properties are rendered in color instead of grayscale. All image channels being used in training for a particular model were then stacked into a single multi-channel image. This input image was then paired with a “label image” from one rater’s visual area or iso-eccentric region boundaries, and together the input and label images constituted a single training example. The correct visual area or iso-eccentric region labels were rendered into image channels such that each area or region was represented by 1 value in a single image channel that was 0 everywhere else. All orthographic projections were made using the *fsaverage*-aligned FreeSurfer spherical surface with the occipital pole aligned to the center of the image in order to minimize anatomical variability.

### Model Training and Hyperparameter Search

Each model was trained using two different sets of hyperparameters, and our training regime was applied ten independent times to each model and hyperparameter set, producing 20 total trained sets of model weights for each of the 16 final models trained in this project. Of the 20 trained models for each combination of input data and predicted labels, the model with the highest overlap of the predicted and hand-drawn labels, as calculated using the Sørensen-Dice coefficient (Dice, 1945; Sørensen, 1948) averaged across labels in the validation dataset, was selected as the final model. Our training regime consisted of 3 rounds of 10 epochs; during each epoch, the model was shown each of its training examples once in a randomized order. After each epoch, if the validation loss was greater than the initial validation loss, the training from that epoch was discarded and the best model from the previous epoch was used in the next epoch instead. The models were evaluated using a loss function that consisted of a weighted sum of the binary cross entropy (BCE; Good, 1963; Cover and Thomas, 2005) and a “dice loss” based on the Sørensen-Dice coefficient. (Our dice loss is defined as one minus the the average Sørensen-Dice coefficient across labels.) The weight for the BCE loss, along with the initial learning rate, the rate of decay of the learning rate after each epoch, the size of the training batches, and the complexity of the encoder ResNet model were hyperparameters of the training (**Tab. 1**). After the first and second rounds of epochs, the learning rate was also reset to 2/3 and 1/3 of its initial value, respectively. Training was performed in PyTorch (Ansel et al., 2024) using the Adam optimizer (Kingma and Ba, 2014).

To find the best hyperparameter values for training the models, we used a hyperparameter grid search. We identified five hyperparameters to optimize, which are detailed in **Table 1**. A set of initial hyperparameter values that resulted in reasonable models was found by experimentation with the training dataset, then, for each hyperparameter, we chose a small number of values around that value as alternative values, forming a hyperparameter grid. We trained our model with all combinations of these hyperparameter values (**Tab. 1**). Specifically, we used (1) two different ResNet architectures: ResNet18 and ResNet34, which differ in that the ResNet34 has more layers and a deeper structure; (2) five different learning rates; (3) five values for γ, which is the constant rate of exponential decay of the learning rate after each step; (4) three different batch sizes; and (5) three different weightings of the BCE versus Dice loss functions. The BCE weight value is the weight to give the BCE-based loss, with the weight of the dice loss equal to 1 minus the BCE weight. During the second and third epochs of training, the BCE weight is reduced to half of its value then to zero, respectively. In order to find the best model performance, we use the same training dataset and validation set for all grid search models. Although we combined two different loss functions to train our model, we only consider the dice loss of the validation dataset to find the optimal model.

**Table 1.** Hyperparameters explored prior to fitting the final models.

| Hyperparameter Name | Tested Values | Model 1 <sup>a</sup>: Best Value | Model 2 <sup>a</sup>: Best Value | Model 3 <sup>a</sup>: Best Value | Model 4 <sup>a</sup>: Best Value |
| --- | --- | --- | --- | --- | --- |
| Base Model <sup>b</sup> | ResNet-18, ResNet-34 | ResNet-18 | ResNet-18 | ResNet-18 | ResNet-18 |
| Learning Rate <sup>c</sup> | $1.67 \times 10^{-3}$, $2.50 \times 10^{-3}$, $3.75 \times 10^{-3}$, $5.62 \times 10^{-3}$, $8.44 \times 10^{-3}$ | $3.75 \times 10^{-3}$ | $1.67 \times 10^{-3}$ | $5.62 \times 10^{-3}$ | $1.67 \times 10^{-3}$ |
| $\gamma$ <sup>d</sup> | 0.80, 0.85, 0.90, 0.95, 1.00 | 0.90 | 1.00 | 0.90 | 0.95 |
| Batch Size <sup>e</sup> | 2, 4, 6 | 4 | 2 | 4 | 2 |
| BCE weight <sup>f</sup> | 0.50, 0.67, 0.75 | 0.75 | 0.75 | 0.5 | 0.5 |

<sup>a</sup>The four models that were tested with each combination of hyperparameters accepted the following inputs and predicted the following outputs: (1) T1-weighted data / visual areas; (2) all data / visual areas; (3) T1-weighted data / iso-eccentric regions; (4) all data / iso-eccentric regions.
<sup>b</sup>The ResNet model we used as an encoder.
<sup>c</sup>The learning rate of updates to the model's parameters during training; the learning rate decays exponentially after each epoch according to the parameter $\gamma$ . Before the second and final rounds of epochs, the learning rate resets to 2/3 and 1/3 of its initial value, respectively.
<sup>d</sup>The rate of decay of the learning rate value ( $\ell$ ) after each training step ( $\ell_{k+1} = \gamma \ell_k$ ) of each epoch.
<sup>e</sup>The number of training images used in each iteration of model training.
<sup>f</sup>The ratio of the BCE loss to the Dice-Sørensen loss during the first round of epochs. During the following round, half of this value is used, and during the final round, the ratio is always 0.

For each set of hyperparameters, we trained four models with different input and label images: (1) T1-weighted data only / visual areas; (2) all available data / visual areas; (3) T1-weighted data only / iso-eccentric regions; (4) all available data / iso-eccentric regions. For each model, we chose the set of hyperparameters that achieved the best loss on the validation dataset (**Tab. 1**, right columns); if multiple sets of hyperparameters were equally effective, we preferred to choose the simpler base encoding model but otherwise chose one of the equally effective hyperparameter sets at random.

When applying these sets of found hyperparameters to the 16 different models with different sets of input and label images, we used the two sets of hyperparameters from the grid search that were associated with the model whose predicted label images were the same; for example when training the model that uses as input T1-weighted data and diffusion-weighted data and that predicts the visual areas as output, we used the hyperparameter set derived from model 1 (T1-weighted data / visual areas) and model 2 (all data / visual areas). All hyperparameter results can be found in **Supplementary Figures 1** and **2**. The final sets of parameters chosen are given in **Table 1**.

Ideally, we would have performed hyperparameter searches for each model being trained, and we would have been exhaustive in evaluating the space of hyperparameters. Such diligence is prohibitively computationally expensive, however, so we chose a strategic subset of the models we would have ideally trained and extrapolated from their results as best we could. Our search included the training of 1,800 models, each trained twice, that varied in terms of input images, the type of boundaries being predicted, and their hyperparameters.

### Generalization and Transfer Learning to a Different Dataset

Because we did not reserve a final validation set of subjects in our model training (see also *AI Ethics*, below) and in order to show that our model has good generalization to other independent datasets, we tested the performance of the final models on an independent dataset collected by different researchers using a different MRI machine: the NYU retinotopy dataset (Himmelberg et al., 2021). Like the HCP dataset, this dataset includes fMRI images from 44 subjects (one subject was excluded due to missing data) including T1-weighted images and retinotopic mapping (BOLD) images. Details are provided elsewhere (Himmelberg et al., 2021), but briefly, subjects participated in sessions that included a T1-weighted scan (TR = 2.4 s; TE = 2.4 ms; voxel size = 0.8 mm^3^ isotropic) and at least one retinotopic mapping scan (TR = 1 s; TE = 37 ms; voxel size = 2 mm^3^) at 3 Tesla using a 3T Siemens MAGNETOM Prisma MRI scanner. Visual stimuli consisted of 192 seconds of drifting bars while the subject fixated at the center of the screen. Data were preprocessed using fMRIprep (Gorgolewski et al., 2011; Esteban et al., 2019), and PRF models were solved using VistaSoft (https://vistalab.stanford.edu/software/, Vista Lab, Stanford University; GitHub: vistalab/vistasoft). Boundaries for V1, V2, and V3 were drawn by at least one researcher on all subjects, but the dataset does not include iso-eccentricity contours.

We tested the performance of the models trained using the HCP dataset on the NYU retinotopy dataset. Because only visual area boundaries and not iso-eccentric boundaries were available for the NYU dataset, we tested only two of the trained CNNs: (1) the CNN that used data from a T1-weighted scan only to predict visual area boundaries and (2) the CNN that used data from a T1-weighted scan and functional (pRF) data to predict visual area boundaries. Because the retinotopic mapping data for the NYU retinotopy dataset is substantially different from that of the HCP dataset (for example, the size and type of the stimulus, the MRI machine and resolution, and duration of the scans all differ between the datasets), we also used our trained model weights as pre-training weights to retrain the CNN that uses T1-weighted and function (PRF) data on a training subset of subjects (*n* = 34) from this new dataset, then test it on a corresponding validation subset (*n* = 9) of the dataset.

### Code and Data Availability

All data used in this project was obtained from previously published sources that are already freely available. Specifically, our primary training dataset uses structural and diffusion MRI data from the WU-Minn Young Adult HCP (Van Essen et al., 2013; https://db.humanconnectome.org/) and functional data from the HCP 7 Tesla Retinotopy dataset (Benson et al., 2018; DOI:10.17605/OSF.IO/BW9EC). Our secondary dataset for testing validation, generalizability, and transfer learning was the NYU Retinotopy Dataset (Himmelberg et al., 2021; DOI:10.18112/openneuro.ds003787.v1.0.1). We have also published a set of derivative data that were processed from these source datasets along with the model weights that are examined in this report (DOI:10.17605/OSF.IO/C49DV). All code used in this project is freely available on GitHub from the repository noahbenson/visual-autolabel. The software environments in which we performed the hyperparameter search and in which we performed training and analyses for this report have been preserved in containerized virtual machines (Docker images: https://docker.com/) and have also been made publicly available (DOI:10.5281/zenodo.14502583).

### AI Ethics

We used cross-validation to separate the whole dataset into a training subdataset (130 subjects) and a validation subdataset (33 subjects). Because the HCP dataset contains a large number of twins, and because twins may have very similar brain structures and retinotopic maps, we required that, if a subject had a relative in the dataset, they and their relative always be part of the same subdataset (training or validation). Because this dataset is small compared to datasets typically used to train a U-net CNN, we did not hold a separate test dataset for post-training validation. Instead, to evaluate whether we overfit the HCP dataset, we tested our final models against an independent dataset, the NYU Retinotopy dataset (Himmelberg et al., 2021), which contains similar data (T1-weighted images and pRF mapping) but was not involved in any training. No changes were made to our our training regime or our final trained models after evaluation against the NYU Retinotopy dataset.

## Results

We trained a total of 16 CNN models to predict either the visual area boundaries of V1, V2, and V3 or the orthogonal iso-eccentric regions from 0–0.5°, 0.5–1°, 1–2°, 2–4°, and 4–7° using various input data (**Fig. 2**) and the set of hyperparameters that was found to perform best for that model.

### CNNs are Highly Accurate at Predicting Visual Area Boundaries

Across the board, models trained to predict either visual area boundaries or quasi-iso-eccentric regions obtained a validation accuracy at or near that of human raters as long as their inputs include functional data—i.e., the same data used by the human raters to draw the boundaries in the first place. **Figure 3** shows this result for the CNNs designed to predict V1, V2, and V3 boundaries. As can be seen in the top panel (**Fig. 3A**), the accuracy of traditional models (“prior” and “inferred”) is consistently below that of any of the CNN models. The “prior” model in this case refers to the anatomical template of retinotopy (Benson et al., 2014; Benson and Winawer, 2018), which is based purely on the anatomical data derived from a T1-weighted image (**Fig. 3C**) and thus is similar in terms of input data to the CNN that received only data derived from a T1-weighted image (**Fig. 3D**). The “inferred” model refers to the result of Bayesian inference applied to both the anatomical (T1-weighted) and functional data combined (Benson and Winawer, 2018) and thus is similar in terms of input data to the CNN that received data from both a T1-weighted image and functional images (**Fig. 3E**). Among CNN models, all models that were trained using functional data as an input (**Fig. 3A**, white background) outperformed any model that did not receive functional data as an input (**Fig. 3A**, gray background). The other two variable types of input data (T2-weighted data and diffusion-weighted data) did not have a substantial effect on the results of the training. All of these models can be compared to the inter-rater reliability of the boundaries (**Fig. 3A**, cyan background; **Fig. 3B**), which serves as a performance ceiling for the trained models. Results for each visual area are provided separately in **Supplementary Figure 3**.

**Figure 3.**
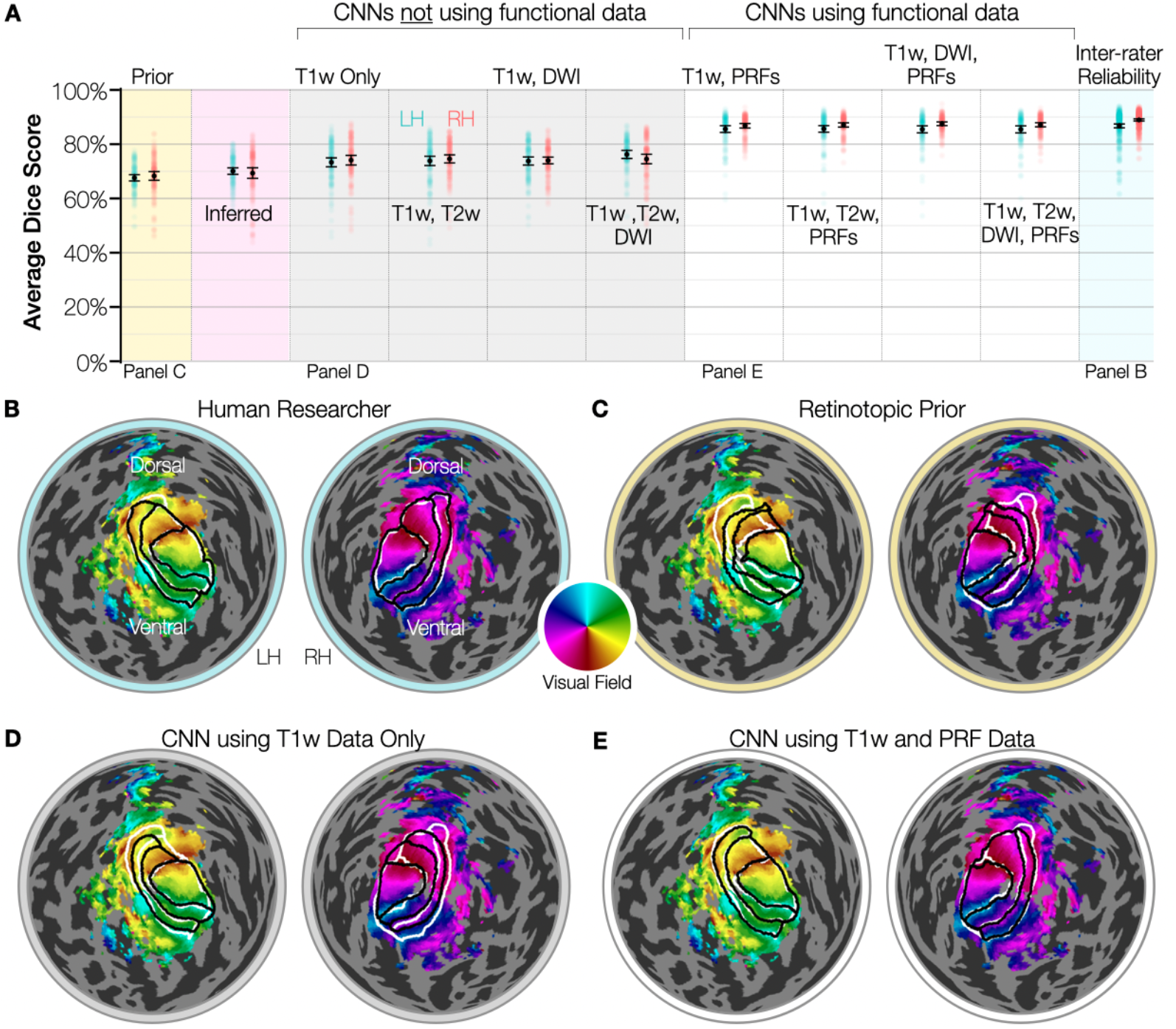
(**A**) The accuracy of each model, as measured using the Sørensen–Dice coefficient averaged over V1, V2, and V3. Points are plotted with partial transparency to show density; each point represents the Dice coefficient between the areas drawn by a single annotator and the areas predicted by the model for a subject in the validation dataset. LH (cyan) and RH (red) hemispheres are plotted separately. Mean values and standard errors are shown in black. The rightmost plot shows the agreement between annotators. The two leftmost plots show the accuracy of the predictions based on FreeSurfer alignment (i.e., the retinotopic prior; yellow background) and based on the Bayesian inference method (magenta background). All models whose accuracies are plotted over a gray background are CNNs that predict V1–V3 boundaries without using functional data, while those with white backgrounds are CNNs that do use functional data. Below panel A are four plots of polar angle on the flattened cortical surfaces of the LH and RH of subject `wlsubj001` from the validation dataset. On each surface, the V1–V3 boundaries drawn by anatomist #1 are shown in white (the white contours are identical in each panel). The black contours are different in each panel and show the predictions made by (**B**) V1–V3 annotator #2, (**C**) the retinotopic prior, (**D**) a CNN trained using only data derived from T1-weighted images, and (**E**) a CNN trained using both T1-weighted data and functional data (i.e. the retinotopic maps).

### CNNs are Highly Accurate at Predicting Iso-Eccentric Regions

A qualitatively similar pattern of results can be found for the prediction of quasi-iso-eccentric regions in **Figure 4**. These results show a few differences from those of **Figure 3**. First, across the board, the Dice accuracy of predicting the quasi-iso-eccentric regions is lower than that for the visual area boundaries, including for the inter-rater reliability. This is at least partly due to the fact that there are five somewhat smaller quasi-iso-eccentric regions and only three somewhat larger visual areas. However, past research has also suggested that eccentricity maps are less conserved across individuals than polar angle maps when one accounts for anatomical variability (Benson and Winawer, 2018), so the reduction in accuracy may also reflect an increased variability in the ground truth. Second, the inferred model (**Fig. 4A**, magenta background) has an accuracy slightly higher than that of the CNNs trained without functional data (**Fig. 4A**, gray background), which was not seen with the visual area predictions in **Figure 3**. This indicates that the Bayesian inference method likely excels at modeling the eccentricity map relative to the polar angle map, which is also supported by the elevated accuracy of the inferred eccentricity maps over the prior eccentricity maps (**Figs. 4A**, yellow background, **4C**). The fact that the inferred map outperforms the CNNs trained without functional data is not surprising given that the inferred method uses functional data as an input. These differences aside, the results largely resemble those of **Figure 3**, with the CNNs that use functional data as input performing at or near the inter-rater reliability. As when predicting the visual area boundaries, the inclusion of T2-weighted data and diffusion-weighted data did not substantially affect the accuracy of the models. Results for each iso-eccentric region are provided separately in **Supplementary Figure 4**.

**Figure 4.**
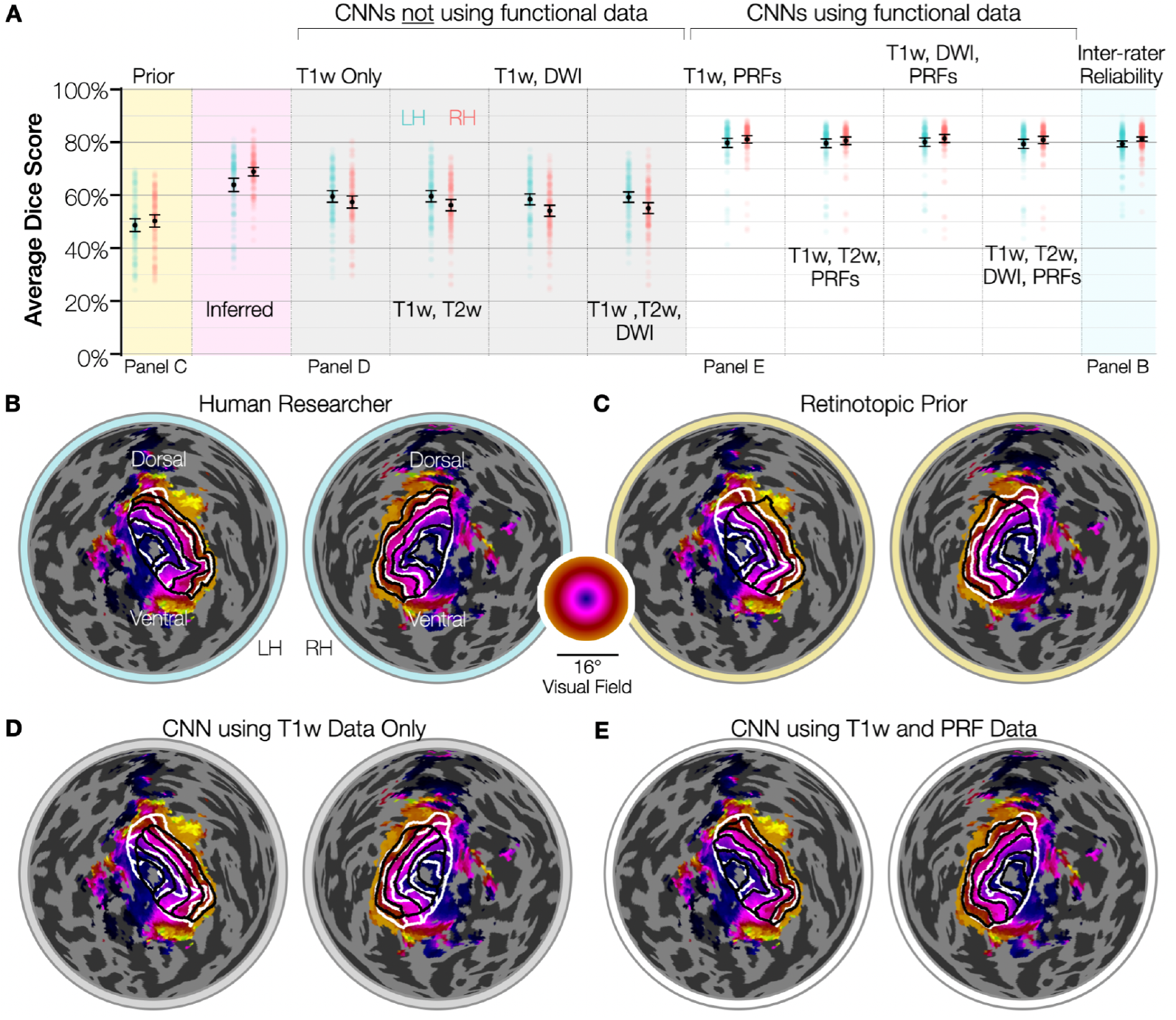
(**A**) The accuracy of each model, as measured using the Sørensen–Dice coefficient averaged over five iso-eccentric regions. Points are plotted with partial transparency to show density; each point represents the Dice coefficient between the areas drawn by a single annotator and the areas predicted by the model for a subject in the validation dataset. Accuracies of LH (cyan) and RH (red) hemispheres are plotted separately. Mean values and standard errors are shown in black. The rightmost plot shows the agreement between annotators. The two leftmost plots show the accuracy of the predictions based on FreeSurfer alignment (i.e., the retinotopic prior; yellow background) and based on the Bayesian inference method (magenta background). All models whose accuracies are plotted over a gray background are CNNs that predict V1–V3 boundaries without using functional data, while those with white backgrounds are CNNs that do use functional data. Below panel A are four plots of polar angle on the flattened cortical surfaces of the LH and RH of subject 115017 from the validation dataset. On each surface, the V1–V3 boundaries drawn by V1–V3 anatomist #1 are shown in white (the white contours are identical in each panel). The black contours are different in each panel and show the predictions made by (**B**) V1–V3 annotator #2, (**C**) the retinotopic prior, (**D**) a CNN trained using only data derived from T1-weighted images, and (**E**) a CNN trained using both T1-weighted data and functional data (i.e. the retinotopic maps).

### Functional Measurements Substantially Improve Accuracy, but Estimated Myelin and Endpoint Maps Do Not

The CNNs trained to predict visual area boundaries can be clustered into two clear groups based on their prediction accuracy: those that used functional data as one of the inputs (**Fig. 3A**, white background) and those that did not (**Fig. 3A**, gray background). All CNNs that used functional inputs achieved similar prediction accuracies, and all CNNs that did not use functional inputs achieved similar prediction accuracies that were uniformly lower than those of that did use functional inputs. The addition of the estimated myelin data and the tract endpoint data did not have a substantial effect on the accuracy of any CNN that we trained. A similar pattern of results can be seen in **Figure 4A**, where performance is also clearly affected by whether or not the model uses functional data as an input, with models that do use functional inputs achieving uniformly higher prediction accuracies than those that do not use functional inputs.

### CNNs Trained on the HCP Dataset Generalize Well to the NYU Dataset

To assess generalizability of our CNN model and to ensure that it did not overfit the HCP dataset, we tested our trained models on the NYU Retinotopy Dataset (Himmelberg et al., 2021), which was not used during training. The NYU Retinotopy Dataset did not include iso-eccentricity contours drawn by human raters, the models were only evaluated against the visual area boundaries. **Figure 5** shows this Dice coefficient when our trained CNNs are used to predict V1, V2, and V3 boundaries in this dataset. When using T1-weighted images as inputs to our trained model, the prediction accuracy closely approximates the prediction accuracy achieved on the HCP dataset (**Figs. 5A**, gray background; **5B**). This demonstrates that our models are not overfitted and that they generalize well to T1-weighted images that were collected for a different study. When using both T1-weighted and functional data with our trained CNNs, the performance is relatively poor. The difference in performance between the CNN trained to use T1-weighted images only and the CNN trained to use both T1-weighted and functional inputs is stark, but can be understood in terms of data variability. T1-weighted images are standard measurements, and FreeSurfer (https://surfer.nmr.mgh.harvard.edu/), which is used to normalize and process the T1-weighted images for our models, is capable of producing similar structural outputs despite noise in the images. Retinotopic maps, on the other hand, depend strongly on many experimental and analysis parameters, such as the size of the stimulus, the kind of stimulus used, and the hemodynamic response function (Lerma-Usabiaga et al., 2020). However, we hypothesize that while the retinotopic maps in the two datasets are substantially different, there is enough similarity in the maps that a small amount of additional training using the NYU dataset would result in a quick recovery of the model accuracy. This recovery should occur quickly because the model is not relearning the entire structure of retinotopic maps but rather is learning the differences between the maps in the two datasets. When we executed this plan to retrain our model with the relatively small NYU dataset (34 training subjects, 9 validation subjects), the model performance achieved a higher accuracy despite being trained on so few subjects (**Fig. 5A**, white background; **Fig. 5C**).

**Figure 5.**
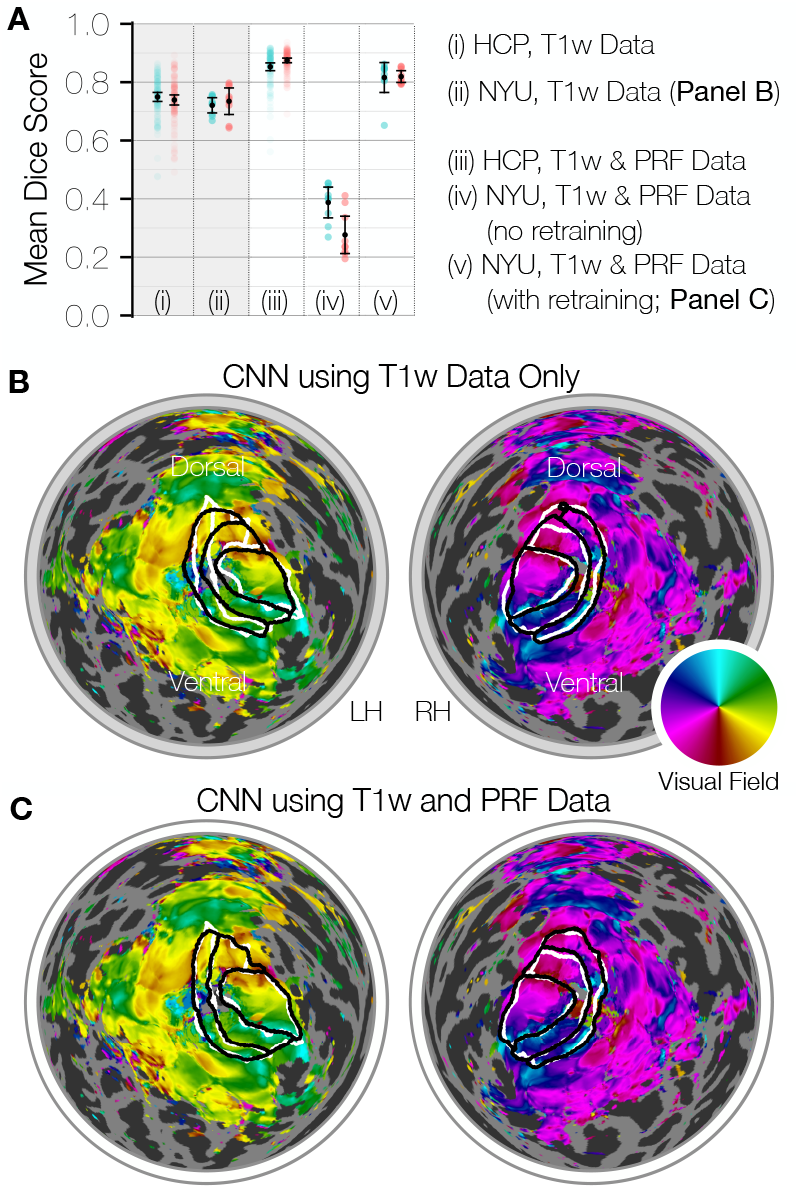
(**A**) The accuracy of each model, as measured using the Sørensen–Dice coefficient averaged over V1, V2, and V3. Points are plotted with partial transparency to show density with plots of the NYU retinotopy dataset appearing as more opaque than those of the HCP dataset due to their lower frequency. Each point represents the Dice coefficient between the areas drawn by a single annotator and the areas predicted by the model for a subject in the validation dataset. Accuracies of LH (cyan) and RH (red) hemispheres are plotted separately. Mean values and standard errors are shown in black. All models whose accuracies are plotted over a gray background are CNNs that predict V1–V3 boundaries without using functional data, while those with white backgrounds are CNNs that do use functional data. Below panel A are two plots of polar angle on the flattened cortical surfaces of the LH and RH of subject wlsubj001 from the NYU validation dataset. On each surface, the hand-drawn V1, V2, and V3 boundaries are shown in white (the white contours are identical in both panels). The black contours are different in each panel and show the predictions made by (**B**) a CNN trained using only data derived from T1-weighted images and (**C**) a CNN trained using both T1-weighted data and functional data (i.e. the retinotopic maps).

## Discussion

### CNNs Can Segment the Visual Cortex with Human Accuracy

This research project follows on many years of work into computational methods for the automatic parcellation of the retinotopic maps in human visual cortex, beginning with the automatic registration methods described by Dougherty et al. (2003). This project establishes for the first time that computational models, specifically CNNs, can achieve a level of performance comparable in accuracy to humans when parcellating functional cortical regions (**Figs. 3** and **4**). The CNNs that achieved this accuracy used, at a minimum, both anatomical (T1-weighted) and functional (pRF) maps to produce human-like parcellations—the same data used by the human raters. This accuracy was achieved not by modeling the visual cortex from first principles but by supervised training from a set of human-drawn parcellation examples.

CNNs that were not trained to use functional maps to make their predictions consistently performed less accurately than those that did use the functional maps. This loss of accuracy can be understood in a number of ways. First, the human raters who parcellated the examples in our training dataset made those parcellations using the functional maps; thus, intuitively, the CNNs most likely to perform well are those that had access to the data that guided the humans. More importantly, however, the drop in accuracy can be understood as a consequence of the observation from previous studies that visual function does not map perfectly onto anatomical structure in the human early visual cortex (Benson and Winawer, 2018). Whether the human raters used the functional maps or any other data to draw the boundaries is less important than whether there is useful information about the boundaries in the data provided to each CNN. If the surface curvature, for example, were tightly coupled to the location of the maps on cortex, then the CNN should in theory be able to learn that coupling when shown surface curvature. The functional maps are de facto tightly coupled to the visual areas because the boundaries are defined in terms of the reversals of the maps, but the anatomical data are coupled to the boundaries only to the extent that the structures they represent are coupled to function.

### CNN Accuracies Provide Limited Evidence of Data Independence

In comparing the performances of different CNNs it is important to remember that CNNs are fundamentally black boxes with no theoretical neuroscientific grounding in which we can interpret them. The fact that one CNN performs better or worse than another CNN does not necessarily mean that the former CNN used input data with a weaker relationship to the predicted quantities. It is possible in such a case that the former CNN was trained using hyperparameters that disadvantaged the processing of its input data relative to that of the latter CNN or that the convolutional structure of the CNNs somehow favored the latter CNN due to some feature of the input data. If a carefully trained CNN matches the inter-rater reliability of a dataset that was not used in training (i.e., achieves human-like accuracy), then that accuracy is strong evidence that there is a strong relationship between whatever input data it is using and the predicted quantities. If, however, another CNN performs less accurately on independent data, that does not necessarily indicate that the relationship between this CNN’s input data and the predicted quantities is weaker.

In the case of the CNNs trained in this project, we cannot conclude with certainty that the input data provided to CNNs that achieved a lower accuracy are fundamentally less useful in solving the problem, only that the CNNs were less effective at solving the problem under our training regime. However, the differences in performance can be considered evidence of such a conclusion, and a number of factors strengthen that evidence. First, we performed a rigorous hyperparameter search (**Tab. 1**; **Figs. S1** and **S2**) in which accuracy was observed to be only weakly sensitive to the choice of hyperparameters, indicating that the problem of predicting boundaries in our dataset is not particularly sensitive to the training regime. Second, our CNNs trained using functional data performed extremely well, indicating that the structure of the CNN we used is well conditioned to solving the problem. Third, our hyperparameter search indicated that the use of a more complex encoder and decoder models (ResNet34 instead of ResNet18) with more internal parameters did not improve model performance, even when using T1-weighted data alone as the input data, suggesting that the models are sufficiently internal complex for the problem. Finally, the accuracies of our models across many training sessions (including the hyperparameter search) are consistent and can be well-predicted by the kind of input data they use, suggesting that they are consistently reaching something like a ceiling in our training.

Beyond the limitations of the method, features of our datasets also substantially limit the reach of our conclusions. Primarily, the functional data used in this project—the HCP 7T Retinotopy dataset and the NYU Retinotopy dataset—contain PRF parameters from one experiment each. The primary training dataset (the HCP), for example, contained measurements from a stimulus with a maximum radius of 8° of eccentricity and whose boundaries were drawn with a maximum of 7° of eccentricity in order to avoid biases in the retinotopic maps that arise near the peripheral boundaries of the stimulus. Our conclusions are thus limited to the mapping between the structure and function of the visual cortex within the foveal 7° of V1, V2, and V3. Any conclusions from the CNNs trained to use data from T1-weighted images alone are arguably less constrained than those from the CNNs using other data because the post-processing of T1-weighted images performed by FreeSurfer is sufficiently mature and thorough that by the time the relevant properties are rendered at the resolution of the training images, small differences arising from imaging parameters, image resolution, or scanner details are minimal.

### CNNs Are More Accurate than Previous Methods when Using Anatomical Data Alone

Anatomical registration methods—i.e., those of or based on FreeSurfer (Fischl et al., 1999)—have been the primary method behind the prediction of visual area organization and boundaries for over a decade (Hinds et al., 2008; Benson et al., 2012; Benson et al., 2014; Benson and Winawer, 2018). The CNNs trained in this project to use information derived only from a T1-weighted image predicted both visual area boundaries and iso-eccentric regions in the validation subjects with accuracies slightly but significantly higher than those of the retinotopic prior of Benson & Winawer (2018). For the first time since the development of these methods, this observation shows that there remains some anatomical information in the posterior visual cortex not already exploited by these registration methods that is predictive of functional organization. Further, the success of our training regime in general suggests, however weakly, that there is little further anatomical information to be exploited.

Interestingly, the CNNs using T1-weighted inputs to predict visual area boundaries (**Fig. 3A**) slightly outperform the Bayesian inference method of Benson & Winawer (2018), which incorporates functional data into its predictions. One reason for this is that the model used by Benson & Winawer did not include any explicit concept of a boundary, and because certain kinds of noise in the retinotopic maps are common near the boundaries such as the blurring of the population responses between the two areas within each voxel, the method was not adept at improving on the boundaries of the retinotopic prior. Rather, the method excelled at aligning the underlying parameter maps, leading to the substantial improvement in the prediction of the eccentricity map (**Fig. 4A**), which Bayesian inference predicts much more accurately than the CNNs that use T1-weighted images alone. In fact, all CNNs trained without functional data perform with a lower accuracy and a higher accuracy variance when predicting the iso-eccentric regions than when predicting the visual area boundaries. This can be partly understood by the larger number of smaller regions being predicted (5 iso-eccentric regions versus 3 visual areas), but also suggests that the eccentricity maps and cortical magnification functions of individuals vary substantially, a finding previously suggested by the range of cortical magnification functions found in the HCP 7 Tesla Retinotopy dataset (Benson et al., 2022).

### Eccentricity Is Less Coupled to Structure than Polar Angle

The distributions of prediction accuracy for CNNs using non-functional data to predict visual area boundaries compared to those using the same data to predict iso-eccentric regions are strikingly different. The accuracies of the CNNs predicting iso-eccentric regions (**Fig. 4A**, gray background) are consistently and substantially both lower and those of the CNNs predicting boundaries (**Fig. 3A**, gray background) and of greater variance. These data are consistent with the observation that the eccentricity maps of subjects contain more variance independent of anatomical structure than the polar angle maps. This theory is similarly supported by the accuracy data from the two registration methods: predictions made using anatomical registration (Benson et al., 2014; Benson and Winawer, 2018) and predictions using Bayesian inference (i.e., functional registration; Benson and Winawer, 2018). Of these two, the method that controls for functional differences has a substantially higher accuracy than the method that controls for anatomical differences when predicting eccentricity-based boundaries (**Fig. 4A**, magenta and yellow backgrounds) than when predicting polar angle-based boundaries (**Fig. 3A**, magenta and yellow backgrounds).

The high variance between subjects in the mapping of eccentricity to structure heavily implies that the cortical magnification function (in terms of eccentricity) varies substantially between subjects. A common simplifying assumption about the retinotopic organization of V1 is that its cortical magnification follows the same function, up to a scaling factor dependent on the individual subject’s V1 surface area, such as that of Horton & Hoyt (1991): *m*(*r*) = *a* / (*r*_0_ + *r*)^2^ where *m* is the cortical magnification in units of mm^2^/deg^2^, *r* is the eccentricity in degrees, *r*_0_ is an offset in degrees that was measured by Horton & Hoyt to be on average 0.75°, and *a* is the constant, in mm^2^, that is a function of the individual subject’s V1 surface area, which was measured by Horton & Hoyt to be on average (17.3 mm)^2^. Horton & Hoyt’s model of cortical magnification has held up exceptionally well in as a prediction of the average magnification in recent studies (Benson et al., 2022), but the simplifying assumption—that the parameter *r*_0_ in this equation is constant across subjects—has not. The observations here are supported by the previous observation that cortical magnification varied substantially in the subjects of the HCP 7 Tesla Retinotopy dataset (Benson et al., 2022).

### The Relationship between Structure and Function Is Closer than Previously Estimated

Previous attempts to quantify the extent to which gray-matter structure is predictive of visual function compared the registrations of subjects to an anatomical average (diffeomorphic structural normalization) to registrations of subjects to a model of retinotopic organization (diffeomorphic functional normalization; Benson and Winawer, 2018). In this prior study, the two registrations were found to be approximately equal and orthogonal, implying that there is approximately as much variance remaining in the functional organization of the retinotopic maps after anatomical contributions to the variance in the maps are removed as was removed from the maps by the anatomical organization.

Here, we have an alternative method of estimating the amount of variance unique to anatomical structure. The CNNs we have trained are clearly more able to exploit anatomical information from gray-matter than registration methods alone are, providing the current best available estimate of the variance in functional maps explainable by anatomy. Simultaneously, the inter-rater reliability puts a ceiling on the accuracy of our gold-standard (**Figs. 3A & 4A**, cyan backgrounds). The performance of the model predicting polar angle-based boundaries from T1-weighted input data is ∼86% the inter-rater reliability while that of the model predicting eccentricity from T1-weighted input data is ∼74% of the inter-rater reliability. This suggests a the variance explained by anatomical structure is much more than half of the variance not explained by structure, as previously estimated.

### T2-weighted Data and Endpoint Maps Failed to Improve CNN Predictions

There was no substantial differences in the performance of CNNs that were trained to use a particular set of data and those that were trained to use that same set of data as well as estimated myelin maps and/or maps of the OR and VOF tract endpoints. Because T2-weighted images and dMR images are substantially easier to collect for most researchers than PRF maps, an improvement in prediction accuracy by CNNs using either of these data sources compared to the accuracy of CNNs using T1-weighted images alone would have been very useful for the field. Unfortunately, in the case of V1, V2, and V3, these additional maps do not appear to provide additional information to the CNNs beyond what is already available in the T1-weighted image data.

Intuitively, the anatomical structure inherent in the white-matter tracts should provide very high quality information about the location and organization of each visual area, assuming that our measurements were of high enough quality and resolution. There are two possible reasons why these maps do not contribute to the improvement of the prediction of boundaries. First, V1 is a region where cortical folding is strongly coupled with area boundaries compared to other cortical regions (Fischl et al., 2008). While connectivity and gray matter microstructure may also be other important anatomical properties characterizing these areas, the utility of these additional maps may be limited by the fact that information from T1-weighted images (including cortical curvature) is already rich enough to enable prediction of the retinotopy of V1 and neighboring maps. In other words, it is possible that these two additional maps can be more useful for predicting areas beyond V3, such as hV4 and V3A. Second, both endpoint maps and estimated myelin maps have limitations regarding their sensitivity and specificity (Reveley et al., 2015; Sandrone et al., 2023). Improved dMRI data acquisition or inclusion of quantitative MRI dataset in future studies may benefit improvement in predicting boundaries.

### The Models Generalize Well to an Independent Dataset

Because our supervised dataset was small compared to those used in typical segmentation problems, we made only a train/test cross-validation split and did not hold out an additional subset of the subjects for final model validation. Instead, we used an independent dataset that contained similar measurements, the NYU Retinotopy dataset (Himmelberg et al., 2021). Because we did not alter our training regime nor retrain our original models after evaluation against the NYU Retinotopy dataset, their performance on this dataset is strong evidence of whether the models are or are not overfit to the particular dataset.

In the case of the models trained to use T1-weighted images, model accuracy is only slightly lower for the NYU dateset than for the HCP dataset. This small difference is likely due to the fact that the peripheral boundary of the visual areas in the HCP dataset was drawn at 7° of eccentricity while in the NYU Retinotopy dataset it was drawn at 11° of eccentricity. However, due to diminishing cortical magnification in the periphery, this difference accounts for a relatively small slice of the visual areas being predicted and thus the accuracy remains only slightly lower than the model accuracy on the validation subjects of the HCP dataset.

In the case of the models trained to use functional data, the accuracy of the model trained only on the HCP data was very low when predicting the boundaries of the NYU dataset. This indicates that this CNN is overfit to the HCP functional data with respect to the NYU functional data. This conclusion is not surprising given the non-trivial differences between the datasets and the small number of samples in the training dataset. The main differences between the functional maps of the datasets have been documented elsewhere (Himmelberg et al., 2021), but in brief, the NYU maps (1) were collected using a larger stimulus and thus yielding larger visual areas on cortex; (2) used a different set of stimuli without rotating wedges or expanding and contracting rings thus producing lower quality maps in the fovea; (3) had systematically higher pRF sizes and eccentricities at equivalent cortical locations; and (4) were collected using different a different scanner at a lower field strength (3T instead of 7T). Nonetheless, we hypothesized that the convolution weights learned via training on the HCP functional data would contain implicit learning about the structure of the functional maps that would be easily transferrable to a new dataset. We thus retrained the CNN using a subset of training subjects from the NYU dataset and the same training regime as we used with the HCP dataset. Despite this dataset being very small (only 34 training examples), an accuracy comparable with that of the HCP dataset was achieved on the NYU validation subjects after the retraining, indicating that the solution learned for the HCP dataset is likely very transferrable to other datasets with slightly different pRF maps.

The datasets were different in a number of ways, but the derived products of the scans are highly similar for anatomical data due to the FreeSurfer processing stream. T1-weighted images are also very standard measurements. PRF experiments on the other hand can be very different and in particular the NYU maps had a different stimulus size and omitted ring and wedge stimuli, leading to very different maps. Nonetheless, transfer learning from the HCP dataset to the NYU dataset was successful.

### Better and More General Models Will Require Better Datasets

Despite evidence that our CNNs generalize well in the case of anatomical input data, it is still the case that each of our CNNs solved a relatively constrained problem using a specific and uniform set of input data. The problem was constrained in several ways.

First, each subject’s maps were pre-aligned to FreeSurfer’s *fsaverage* atlas surface in order to reduce anatomical variability prior to the production of input or label images. Unlike many other ways in which this problem was constrained, this constraint simplifies the work being done by the models substantially but likely reduces their generalizability only a small amount. This leverage is possible because the *fsaverage* alignment produced by FreeSurfer is quite general for most subjects. Subjects with lesions or with large T1-weighted imaging artifacts, however, are difficult to align to anatomical templates, and thus they are likely to be poorly predicted by our models. In principle, there is no reason that a CNN, possibly one designed to use 3D MR images as inputs and to produce voxel-wise output labels, could not predict functional area boundaries without using FreeSurfer-based post-processing. Such a CNN would be more general and could theoretically be used with lesioned brains. However, because the problem of locating V1–V3 in a T1-weighted image is substantially more difficult than finding it in an anatomically normalized flattened 3D projection, the size of the training data would likely need to be substantially larger than our training dataset to achieve a similar accuracy.

A second constraint on our models’ solutions arises from the heterogeneity of the subjects in the Young Adult HCP dataset. The HCP retinotopy subjects in particular were constrained by good health and age (20–40 years old), had relatively modest racial and ethnic variability, contained a large overrepresentation of twins, and consisted of 60% females and 40% males. This dataset is likely to generalize to more diverse datasets only to the extent that the organizations of V1, V2, and V3 are invariant to dataset features such as age and twin status. Although replication of the results in an independent dataset with an independent subject population is encouraging evidence that the images derived from T1-weighted measurements that were used for our model generalize well, much larger and more diverse retinotopic mapping datasets are needed before this kind of generalizability can be properly measured.

A final constraint on our models comes from the uniformity in scan protocol across subjects. All subjects in the HCP were scanned with a single protocol using well calibrated equipment and a single post-processing pipeline. While many studies contain data of a quality similar to that of the HCP, many studies are not able to devote as much attention or scan time to their experiments and post-processing. In comparing to the NYU retinotopy dataset (**Fig. 5**), we see that the model using T1-weighted data alone as input generalizes well across datasets despite differences in site, scanner, and subject pool, again suggesting that T1-weighted images and the pipelines for processing them generalize well across datasets. Simultaneously, the failure of the model using functional inputs to generalize well to the NYU dataset demonstrates a limit of the models’ generalizability—one that could be overcome with a dataset of diverse retinotopic maps collected and processed using a variety of stimuli, protocols, and software.

### Our Models Can Be Easily Used By Other Researchers

All code and data used in this project are publicly available (see *Methods: Code and Data Availability*). The source code used in this project was written with the intent that other researchers be able to easily adapt it to a variety of different kinds of functional data that might be useful for predicting the boundaries or features of other brain regions, broadly.

Additionally, we have made the trained models freely available (DOI:10.17605/OSF.IO/C49DV), and we have written a command-line tool into the project’s library that allows one to easily and quickly apply any of the models to a novel subject for which the appropriate data has been collected. This tool has been preserved in a containerized virtual machine (see *Methods: Code and Data Availability*), and instructions for using it can be found in the *Supplemental Materials*.

## Acknowledgements

This work was supported by National Eye Institute grant 1R01EY033628 (to N.C.B and J.W), the Japan Society for the Promotion of Science KAKENHI Grants JP21H03789 and JP24K03240 (H.T.), and the National Institute of Information and Communications Technology Project for Collaborative Research in Computational Neuroscience (CRCNS): Innovative Approaches to Science and Engineering Research on Brain Function (H.T.).

