## Supplemental Materials for "Machine Learning Matches Human Performance at Segmenting the Human Visual Cortex"

#### Convolutional Neural Networks can Segment the Human Visual Cortex with Human Accuracy

### Reproducing the Analyses in this Project

The analyses in this project were produced using a software environment that has been preserved in Docker images (https://docker.com/). These images were published prior to this publication and are freely available (DOI:10.5281/zenodo.14502583). The image named `analysis.tar.gz`, in particular, contains an environment capable of reproducing the computations in this project. When this Docker image is run with the argument `jupyter`, it starts a Jupyter server in which the analysis notebook for the project can be executed. The image additionally can be run to apply either of the models trained to use T1-weighted data alone to predict visual areas or iso-eccentric regions to a novel FreeSurfer subject. It can also be invoked using the Docker command-line tool; for example, to apply the model to a FreeSurfer subject stored in the directory `/store/subjects/bert`, use the following command:

```
docker run -it --rm                     \
           -v /store/subjects:/subjects \
           nben/benson2025-unet:latest bert
```

For additional information, see also this command:

```
docker run -it --rm nben/benson2025-unet:latest --help
```

Additional instructions on how to use these models and Docker images as well as all other data produced for this report can be found in the Open Science Framework page for this project (DOI:10.17605/OSF.IO/C49DV). This page includes a wiki along with all endpoint maps, trained models, hyperparameter grid-search results, and training and validation images.

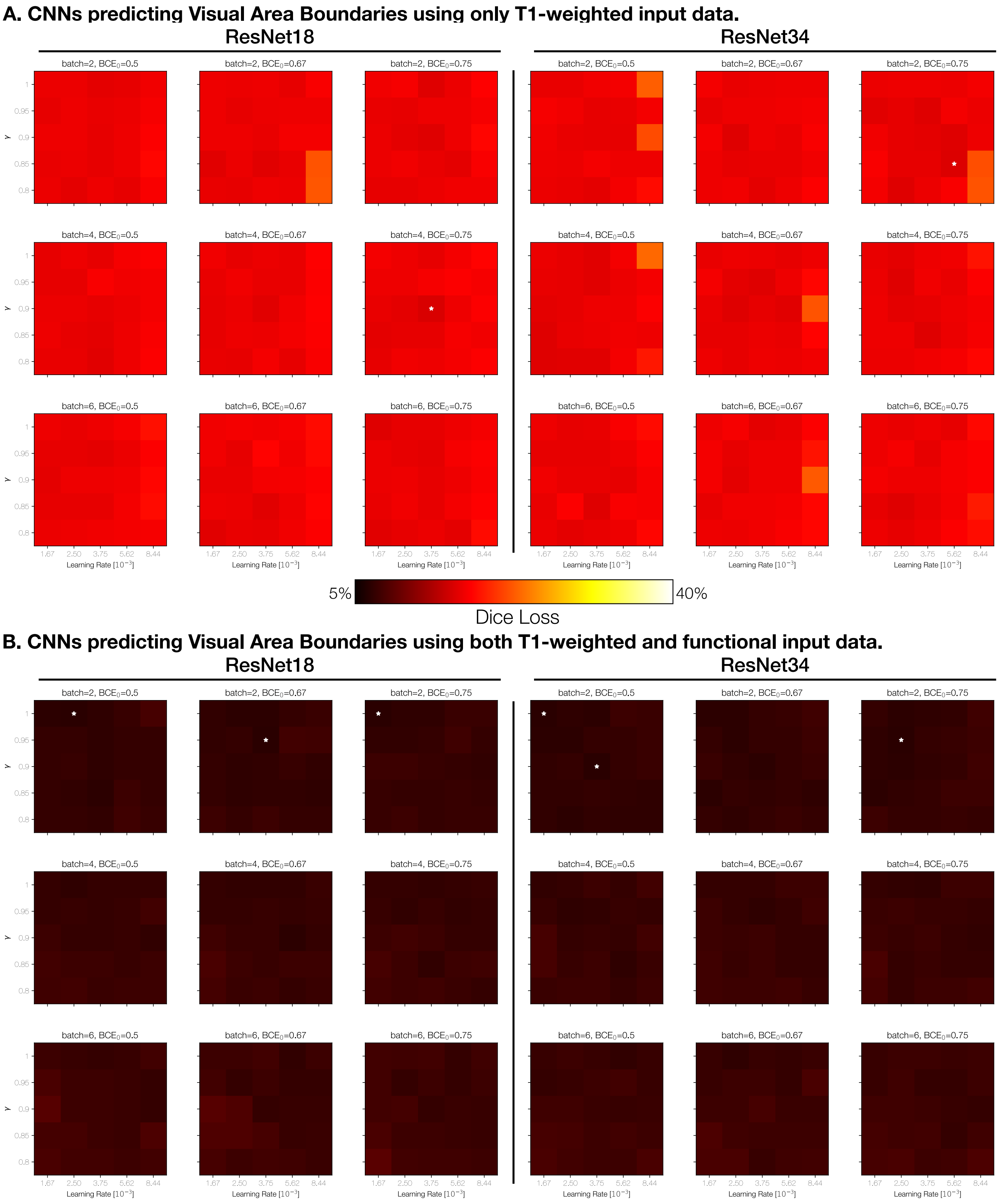


**Supplementary Figure 1**. Complete results of the grid-search for the CNNs that predicted visual area boundaries using (**A**) T1-weighted data alone and (**B**) T1-weighted data and functional data. Results are plotted in terms of the dice loss, with smaller values indicating a higher overlap between predicted and gold-standard areas in the validation dataset. Each cell represents one model with a unique set of inputs and hyperparameters. White stars indicate cells whose performance was within 1% of the best performance across hyperparameters.

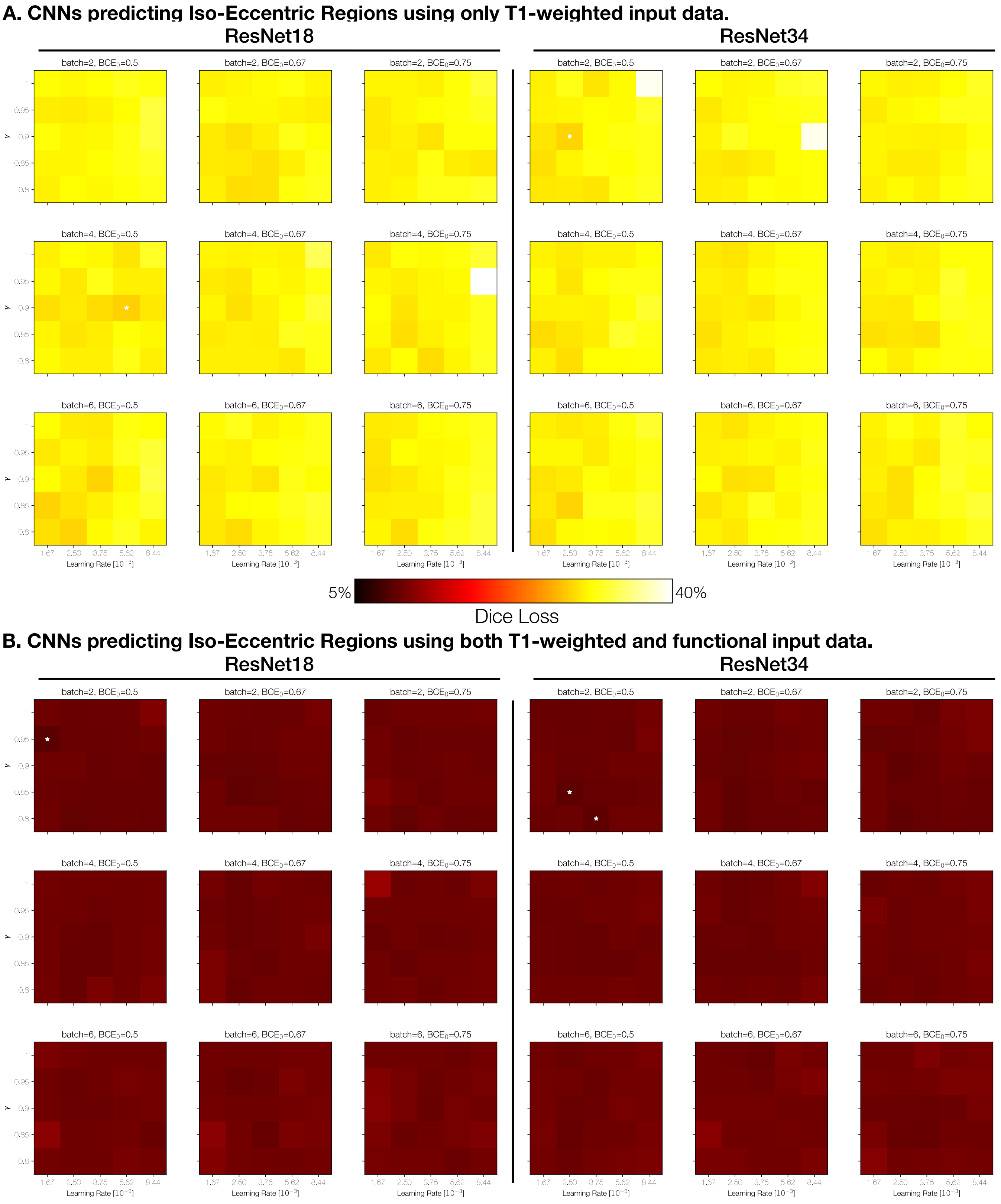


**Supplementary Figure 2**. Complete results of the grid-search for the CNNs that predicted iso-eccentric regions using (**A**) T1-weighted data alone and (**B**) T1-weighted data and functional data. Results are plotted in terms of the dice loss, with smaller values indicating a higher overlap between predicted and gold-standard areas in the validation dataset. Each cell represents one model with a unique set of inputs and hyperparameters. White stars indicate cells whose performance was within 1% of the best performance across hyperparameters.

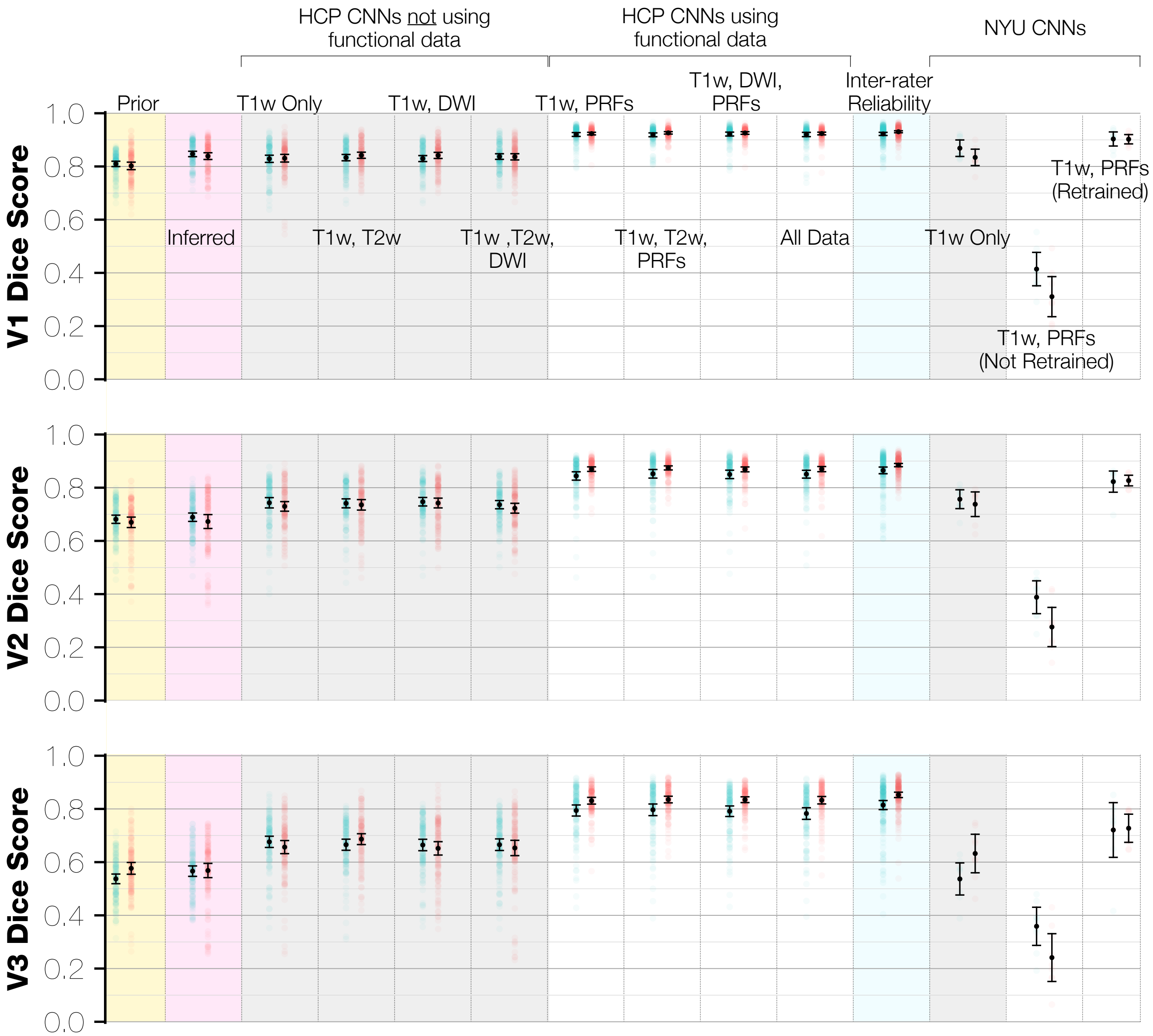


**Supplementary Figure 3**. CNN prediction accuracies for visual area boundaries, separated by each visual area and the input data used to train the model. Background colors in each panel are as in **Figure 3**.

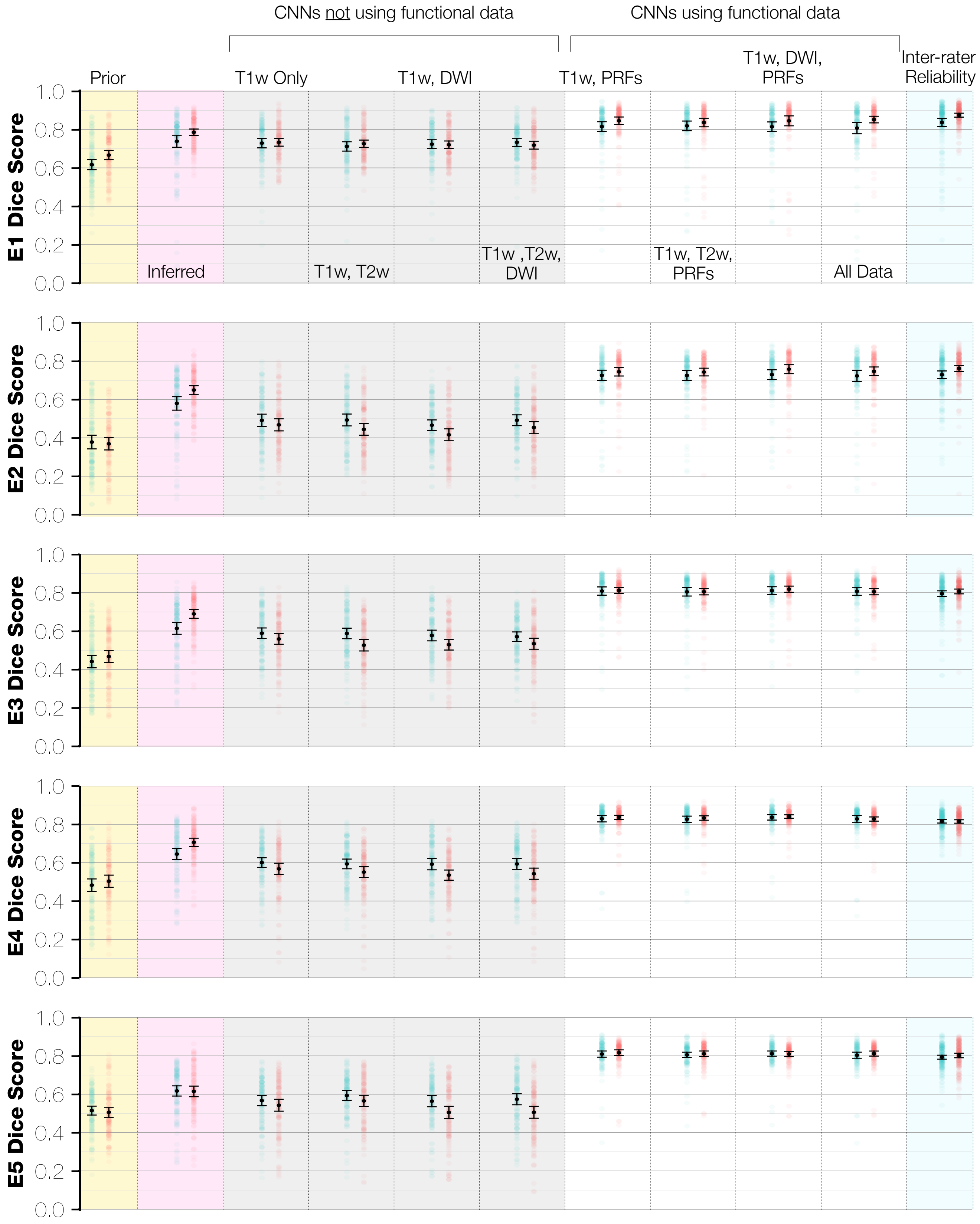


**Supplementary Figure 4**. CNN prediction accuracies for iso-eccentric regions, separated by each region and by the input data used to train the model. Background colors in each panel are as in **Figure 4**. The iso-eccentric regions are 0–0.5° (E1), 0.5–1° (E2), 1–2° (E3), 2–4° (E4), and 4–7° (E5).
